# Probing adaptation and spontaneous firing in human auditory-nerve fibers with far-field peri-stimulus time responses

**DOI:** 10.1101/2021.03.08.434366

**Authors:** Antoine Huet, Charlène Batrel, Xavier Dubernard, Jean-Charles Kleiber, Gilles Desmadryl, Frédéric Venail, M. Charles Liberman, Régis Nouvian, Jean-Luc Puel, Jérôme Bourien

**Author notes:** These authors contributed equally to this work (Antoine Huet, Charlène Batrel, Jean-Luc Puel, Jérôme Bourien).

## Abstract

Information in sound stimuli is conveyed from sensory hair cells to the cochlear nuclei by the firing of auditory nerve fibers (ANFs). For obvious ethical reasons, single unit recordings from the cochlear nerve have never been performed in human, thus functional hallmarks of ANFs are unknown. By filtering and rectifying the electrical signal recorded at the round window of gerbil cochleae, we reconstructed a peri-stimulus time response (PSTR), with a waveform similar to the peri-stimulus time histograms (PSTHs) recorded from single ANFs. Pair-by-pair analysis of simultaneous PSTR and PSTH recordings in gerbil provided a model to predict the rapid adaptation and spontaneous discharge rates (SR) in a population of ANFs according to their location in the cochlea. We then probed the model in the mouse, in which the SR-based distribution of ANFs differs from the gerbil. We show that the PSTR-based predictions of the rapid adaptation time constant and mean SR across frequency again matched those obtained by recordings from single ANFs. Using PSTR recorded from the human cochlear nerve in 8 normal-hearing patients who underwent cerebellopontine angle surgeries for a functional cranial-nerve disorders (trigeminal neuralgia or hemifacial spasm), we predicted a rapid adaptation of about 3 milliseconds and a mean SR of 23 spikes/s in the 4 kHz frequency range in human ANFs. Together, our results support the use of PSTR as a promising diagnostic tool to map the auditory nerve in humans, thus opening new avenues to better understanding neuropathies, tinnitus, and hyperacusis.

## Introduction

Hearing relies on auditory nerve fibers (ANFs), which convey the neural spike trains initiated by the sensory cells of the cochlea to the cochlear nuclei. To achieve intensity coding over a large dynamic range, ANFs differ in their threshold of activation and saturation. In addition, the mapping of the ANFs across the tonotopic axis of the cochlea allows the best frequency coding consistent with the hearing sensitivity (Meyer et al., 2009) and ecological functions of the species (Taberner and Liberman, 2005; Huet et al., 2016). Beside their spontaneous spike rate and threshold for activation, their evoked stereotyped response is also a physiological hallmark of ANFs. At the onset of sound-stimulation, the action potential firing rate first increases and reaches an onset peak. The spike rate then declines monotonically to a steady-state value (plateau). Upon cessation of the stimulus, the spike rate drops below the spontaneous rate before gradually recovering (Heil and Peterson, 2015).

Given their role in sound coding, characterizing ANFs populations and their properties is an important approach to studying auditory deficits. However, the compound action potential (CAP) of the auditory nerve and the wave I of the auditory brainstem response only capture the onset responses of ANFs, and assessment of other aspects requires invasive and difficult techniques, such as inserting a glass microelectrode into the auditory nerve (Bourien et al., 2014). An alternative approach is to extract the ANFs’ temporal responses from round-window recordings using electrocochleography (ECochG). This recording method is currently used in animals and humans to extract both the cochlear microphonic and the summating potentials reflecting the summed hair-cell receptor potentials, and the compound action potential (CAP) that represents the summed action potentials firing in synchrony at the onset of the acoustic stimulus (Eggermont, 2017). In response to narrowband noise, the CAP is followed by small variations related to stimulus envelope fluctuations (Joris et al., 2004). After filtering and rectifying, the electrical signal recorded at the round window remarkably replicates the shape of peri-stimulus time histograms (PSTHs) of single ANFs (Cazals and Huang, 1996), and thus corresponds to what can be expected from composite PSTHs of the whole nerve (Kiang, 1990). Here, we investigated the composite PSTH as peri-stimulus time responses, referred as PSTRs. Simultaneous recordings show that PSTR displays comparable kinetics of adaptation to those measured from PSTHs derived from single units. Additionally, PSTR nicely predicts the rapid time constant and the spontaneous rate (SR) firing of ANFs in gerbils and mice. Finally, we provide a proof of concept, supporting the use of PSTRs in humans as a promising diagnostic tool to better understand auditory nerve dysfunctions such as neuropathies, tinnitus, and hyperacusis.

## Materials and methods

### Gerbil and mouse experiments

Young adult Mongolian gerbils and C57BL/6 strain mice were obtained from Charles River Laboratories (L’Arbresle, France). Animals were housed in facilities accredited by the French ministry of agriculture and forestry (Ministère de l’Agriculture et de la Forêt, Agreement C-34-172-36), and the experimental protocol was approved (Authorization CEEA-LR-12111) by the Animal Ethics Committee of Languedoc-Roussillon (France). Experiments were carried out in accordance with the animal welfare guidelines 2010/63/EC of the European Communities Council Directive regarding the care and use of animals for experimental procedures. All efforts were made to minimize the number and suffering of the animals used.

### Round-window recordings

Gerbils or mice were anesthetized by an intraperitoneal injection of a mixture of Rompun 2% (3 mg/kg) and Zoletil 50 (40 mg/kg). The left cochlea was exposed though a retroauricular surgical approach. The recording electrode was placed on the bony edge of the round window (RW) membrane of cochlea. The bulla (including the recording electrode) was then closed with dental cement. Electrophysiological recordings were performed in a Faraday shielded, anechoic, sound-proof cage. Animals were placed on a vibration-isolated table (TMC, Peabody, MA, USA). The rectal temperature was measured with a thermistor probe, and maintained at 38 °C ± 1°C using a heated blanket beneath the animal. The acoustic stimuli were delivered under calibrated conditions using a custom acoustic assembly comprising a signal generator (PXI-4461 controlled by LabVIEW, National Instrument Company), an audio amplifier (Tucker Davis, SA1) and a magnetic speaker (Tucker Davis, MF1).

Electrical signals were recorded in response to bursts (trapezoidal envelope, 2.5 ms rise and fall) of frequency band noise using different parameters such as frequency bandwidth, interstimulus time interval, level, and central frequency. Two consecutive bursts (called a pair) were presented in opposite polarity to reduce the microphonic potentials originating from transduction currents in hair cells. Each pair was designed to be mutually independent by refreshing the seed of the pseudorandom noise generator (**Fig. 1a,b**). The signal resulting from the half sum within each pair was 300-1200 Hz filtered and the temporal envelope extracted by applying a full-wave rectification and a smoothing filter with a 1-ms span time (**Fig. 1c, d**). The peri-stimulus time response (PSTR) was obtained by averaging over all pairs (**Fig. 1e**).

**Fig. 1.**
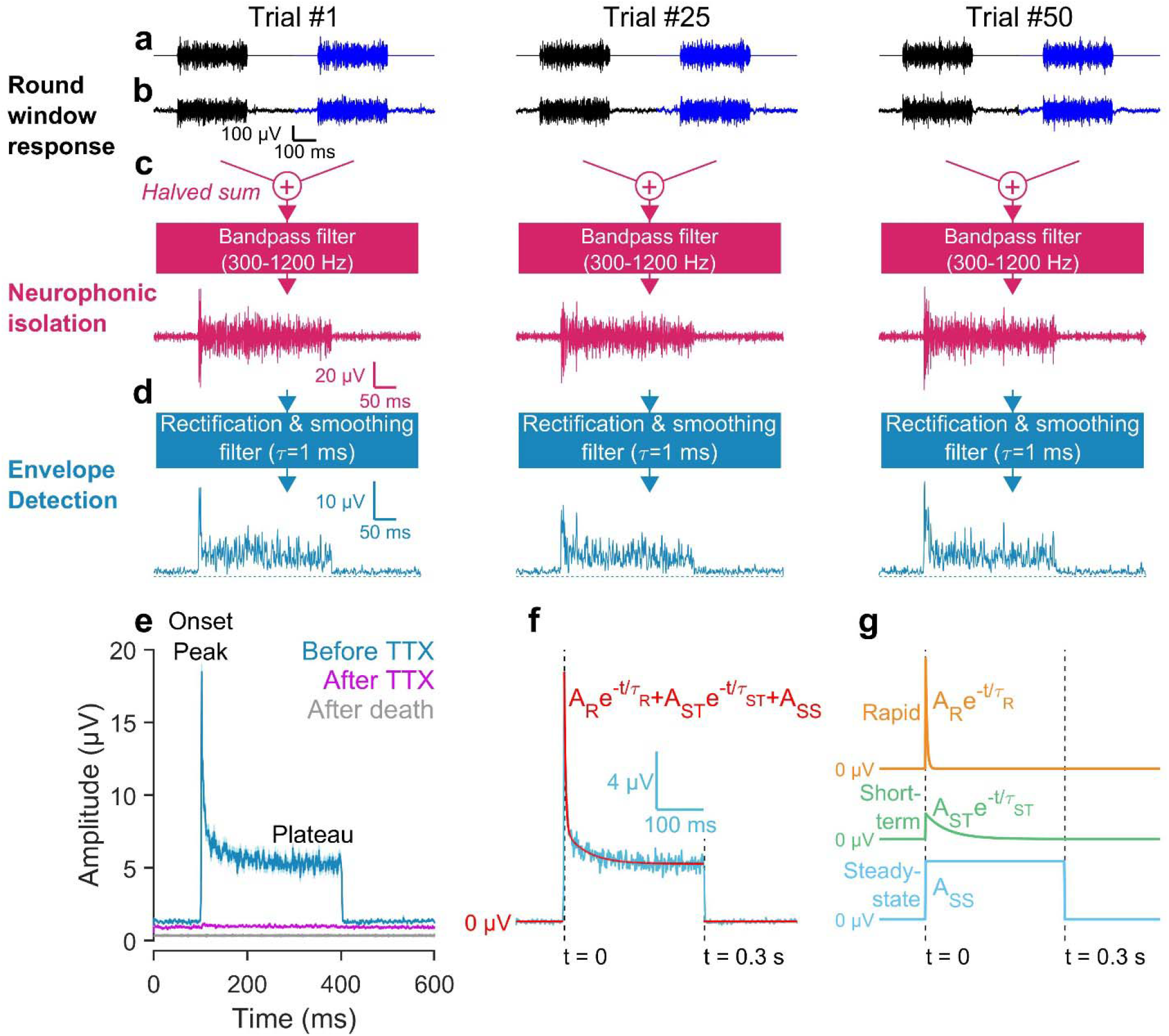
Peri-stimulus time response (PSTR) of the auditory nerve. **a, b** Electrical signal recorded at round window (B) in response to 300-ms bursts of 1/3 octave frequency band noise centered to 4 kHz (A) and presented at 50 dB SPL. **c** Neurophonic isolation by *i*) calculating the halved-sum of individual responses within each pair and *ii*) band-pass filtering to catch the neuronal component generated by the auditory nerve (Batrel et al., 2017). **d** Temporal envelope extraction of the neurophonic potential by applying a full-wave rectification and a smoothing filter (1-ms span time). **e** A PSTR was obtained by averaging the individual traces shown in D. Infusion of 10 µM TTX into the round window niche completely abolished the PSTR. Note the resistance of baseline to TTX (purple trace) as compared to death (gray trace). **f, g** PSTR time course during the sound stimulation fits with a 2 decaying-exponential model (rapid and short-term components) plus a constant term.

### Simultaneous round-window and single auditory nerve fiber recordings

Simultaneous recordings from the RW and the auditory nerve were only performed in gerbils. The round window electrode was positioned as described above. Next, animals were prepared for single-unit recordings from the auditory nerve, as described in Huet *et al* (Huet et al., 2018). Briefly, animals were placed in a custom head holder and their body temperature monitored and maintained at 38 °C ± 1°C. The calibrated acoustic stimuli were delivered to the tympanic membrane through magnetic speakers (type MF1, Tucker-Davis company) coupled to the ear bars. The left cochlear nerve was exposed using a posterior fossa approach. Extracellular action potentials from single auditory nerve fibers were recorded with glass microelectrodes (*in vivo* resistance between 80 and 110 MΩ) connected to an AxoClamp 2B (Molecular Devices), filled with 3 M NaCl. A silver-silver chloride reference wire was placed in the animal’s neck musculature. The fiber’s SR was evaluated by averaging the firing (spikes/s) over 30 s. Fiber CFs were measured using a threshold-tracking program (10 spikes/s > SR). Concomitant recordings were obtained in response to the PSTR stimulus described above, in which the frequency bandwidth was centered on the fiber’s CF.

### Proof of concept in humans

The electrophysiological recordings from the auditory nerve in humans were performed in the Reims University Hospital. Patients underwent microvascular decompression to relieve trigeminal neuralgia (*n* = 7) and hemifacial spasm (*n* = 1) via the retrosigmoid approach (Møller and Jannetta, 1981; Møller and Jho, 1989). The recordings presented in this report were made as part of the routine intraoperative monitoring of auditory evoked potentials. Such monitoring minimizes the risk of hearing loss resulting from manipulation of the eighth nerve. The ethics committee of “*Sud-Méditerrannée*” approved this study (MelAudi-2). All the subjects gave their informed consent to participate to this clinical trial (ClinicalTrials.gov identifier: NCT03552224). The average age of the patients was 62.1 ± 9.3 years, and they had normal or sub-normal auditory thresholds (≤ 20 dB HL) between 500 and 4000 Hz (**Supp. Fig. 4**). Monitoring was based on compound action potentials (CAP) in responses to clicks varying from 0 to 80 dB SPL. At the end of the decompression procedure, PSTR were recorded in response to bursts of third-octave-band noise (200-ms duration, 2.5 presentations/s, 100 presentations) centered on 4 kHz, and presented 40 dB above the click-evoked CAP threshold.

### Data fitting

Given the shape of PSTR recorded at the cochlea round window, we used the same fitting model as that proposed by Westerman and Smith to characterize the adaptation of the firing rate in auditory nerve fibers (Westerman and Smith, 1984). This fitting model consists of two decaying components (rapid and short-term) plus a constant term: 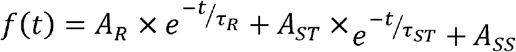 where *A*_*R*_ is the amplitude and τ_*R*_ the time constant of the rapid component, *A*_*ST*_ the amplitude and τ_*ST*_ the time constant of the short-term component, and *A*_*SS*_ the steady-state component reflecting the plateau (**Fig. 1f, g**). The origin of the time coordinate (*t* = 0) in the fitting model is aligned with the PSTR onset-peak latency and the basal activity measured during the 50ms preceding the PSTR onset peak was subtracted from the PSTR to removed non-neural and non-cochlear electrical contributions. The fitting was carried out using a non-linear, least-squares procedure (*fit* function in Matlab) and 6 parameters of the PSTR were considered: *A*_*R*_, *A*_*ST*_, *A*_*SS*_,τ_*R*_,τ_*ST*_ and the PSTR peak-to-plateau ratio 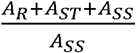.

PSTHs were analysed with the same fitting procedure. However, because of the ANF refractoriness, a silent period frequently appears in the 2-ms following the PSTH onset-peak, making the measurement unreliable. We thus removed this silent period by interpolation between the onset bin and the first bin following the silent interval, as previously used by Westerman and Smith (Westerman and Smith, 1984).

### Statistics

Data are expressed as mean ± SEM. Normality of the variables was tested by the Shapiro-Wilks test. If conditions for a parametric test were met, the significance of the group differences was assessed with a one-way ANOVA; once the significance of the group differences (*p* < 0.05) was established, Tukey’s *post hoc* tests were subsequently used for pairwise comparisons. If conditions were not net, Kruskal–Wallis tests were used to assess the significance of differences among several groups; if the group differences were significant (*p* < 0.05), Dunn’s tests were then used for *post hoc* comparisons between pairs of groups.

## Results

Using electrocochleography in gerbil, we recorded mass potentials at the round window that were evoked by a train of narrow-band noise bursts centered on a probe frequency (see also Batrel *et al*, 2017 (Batrel et al., 2017)). A trial consisted of a pair of bursts with opposite polarities to reduce the cochlear microphonic originating from outer hair cells. The response was then filtered at 300-1200 Hz to isolate the neural component and smoothed with a 1 ms span time fiber (**Fig. 1a-c**). The composite PSTH of the whole nerve (Kiang, 1990; Cazals and Huang, 1996) was built from the average across trials of a full-wave rectified and low-pass filtered electrical signal (**Fig. 1d**). Consistent with its neuronal origin, acute 30-min round-window infusion of 10 μM tetrodotoxin (TTX) completely abolished the response. Note however the TTX resistance of the baseline (1.73 ± 0.09 µV to 1.02 ± 0.05 µV before and after TTX, respectively), and the drop into the noise floor after death (0.30 ± 0.01 µV), suggesting a contribution from some non-neuronal or non-cochlear origin **(Fig. 1e)**. Thus, to quantify the sound-evoked response that we called post-stimulus time response (PSTR), we subtracted the baseline from the evoked response. The PSTR hallmarks consisted of an onset stimulation peak followed by adaptation, plus a plateau until end of the stimulus. Similarly to the PSTH from a single ANF (Westerman and Smith, 1984), the PSTR can be well fit as the sum of two decaying exponential components and a plateau (rapid, 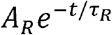; short-term, 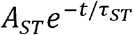; steady-state, ASS; **Fig. 1f, g**). Because of the similarity of shape with the PSTH of a single ANF, and its TTX sensitivity, the PSTR likely reflects the activity of a population of fibers within the auditory nerve.

### Characterization of the peri-stimulus time responses

#### Dependence on the stimulus bandwidth

Passband filtering of the noise bursts produces envelope fluctuations that modulate the temporal response of the auditory nerve (Dau et al., 1999; Joris et al., 2004). Thus, PSTR will be sensitive to the frequency bandwidth of the noise. To determine the most efficient sound stimuli, we compared PSTRs evoked by noise-band bursts of different bandwidths to those evoked by pure tones (representative example in **Fig. 2a, b**). The central frequency of the noise band was fixed at 4 kHz (*i*.*e*. the best sensitivity region of the gerbils’ hearing range (Ryan, 1976; Huet et al., 2016)), and the level set to 50 dB sound pressure level (SPL). While the PSTR onset peak remained constant across all conditions, reduction of the bandwidth led to an amplitude decrease of the plateau, which almost completely disappeared for a pure tone (**Fig. 2a, b**).

**Fig. 2.**
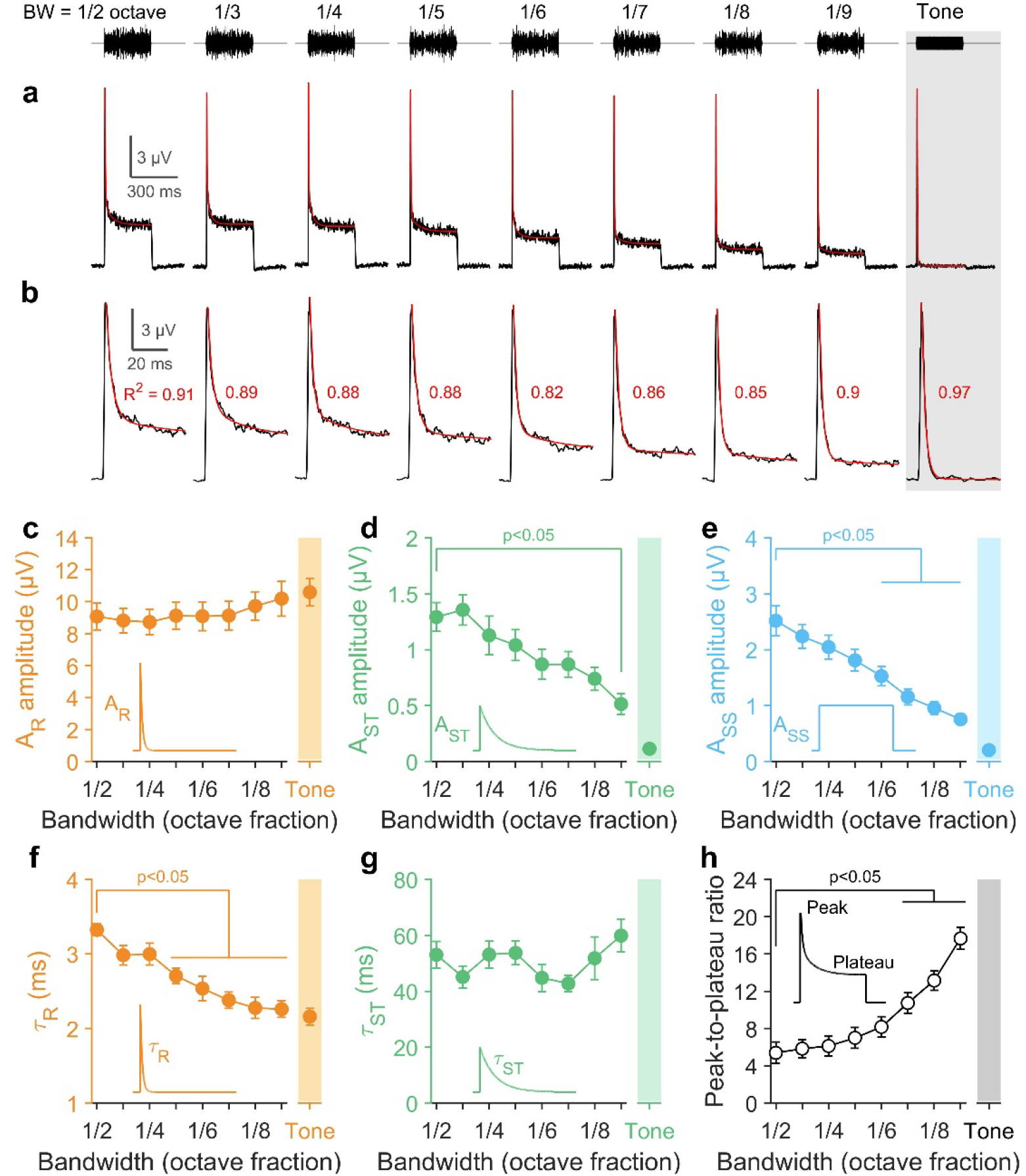
PSTR as a function of stimulus frequency bandwidth expressed as octave fractions. **a** PSTRs (bottom traces) in response to a burst of band-limited noise (top trace) centered at 4 kHz and varying in bandwidth (BW) from 1/2 to 1/9 octave band (data from one representative gerbil). The PSTR obtained to a tone burst presented at 4 kHz is shown on the right (gray panel). The level of presentation was fixed at 50 dB SPL for all stimuli. **b** Magnification of the PSTR onset peak shown above in panel A. In red, the fitting model (*i*.*e*. two decaying-exponential components plus a constant term) and its coefficient of determination (*R*^2^, red digit). **c-e** Amplitude of three components of the fitting model as a function of the bandwidth. **f, g** Time constant of the rapid and short-term components as a function of the stimulus bandwidth. **h** Peak-to-plateau ratio as a function of the bandwidth. Data are mean ± sem, *n* = 12 gerbils. Statistical comparisons between pairs of samples were performed using the 1/2 octave band as the reference. Data obtained in response to a tone burst are shown using a colored back.

A least-squares curve-fitting method was used to quantify the amplitude (**Fig. 2c-e**) and the time constant of the PSTR (**Fig. 2f, g**). The amplitude of the rapid component *A*_R_ was resistant to the bandwidth reduction and showed a slight non-significant increase (9.1 ± 0.8 µV and 10.6 ± 0.9 µV in response to a half-octave bandwidth and pure tones, respectively, **Fig. 2c**). In contrast, the amplitude of the short-term (*A*_ST_) and the steady-state (*A*_SS_) components decreased for smaller bandwidths (from 1.3 ± 0.12 µV to 0.12 ± 0.05 µV and 2.5 ± 0.27 µV to 0.25 ± 0.03 µV for half-octave bandwidth and pure tone, respectively; **Fig. 2d, e**). The kinetics of the rapid component became faster with bandwidth reduction (τ_*R*_: 3.3 ± 0.08 ms to 2.2 ± 0.1 ms for half-octave bandwidth and pure tone, respectively), while the kinetics of the short-term (τ_*ST*_) adaptation component remained insensitive to bandwidth changes (**Fig. 2f, g**). Together, this led to an increase of the PSTR peak-to-plateau ratio from 5.4 ± 0.4 to 17.7 ± 2.5 for 1/2 to 1/9 octave bandwidths, respectively (**Fig. 2h**). Given that the amplitudes and time constants remained constant up to 1/3 bandwidth noise, this bandwidth was used in subsequent experiments.

#### Dependence on the inter-stimulus time interval

The ability of the ANFs to respond consistently to repeated acoustic cues depends strongly on the time between successive stimulations. Short silent intervals decrease the strength of ANF onset responses, because of the depletion of the readily releasable pool of presynaptic vesicles, and the time required to replenish them (Beutner et al., 2001). To evaluate the behavior of the PSTRs in this masking phenomenon, we varied the inter-stimulus intervals between two consecutive 4 kHz bandwidth stimulations from 40 to 600 ms (**Fig. 3a**). Raising the inter-stimulus time interval from 40 to 300 ms increased the onset-peak amplitude. Above 300 ms, the onset peak remained constant. We then expressed the amplitude parameters derived from the fits as the function inter-stimulus interval (**Fig. 3b-d**). Like the onset-peak, the amplitude of the rapid and short-term components increased up to 300 ms inter-stimulus intervals, but did not change beyond 300 ms (*A*_R_: 4.9 ± 0.5 µV to 9.0 ± 0.8 µV and *A*_ST_: 0.5 ± 0.1 µV to 1.5 ± 0.2 µV, from 40 to 300 ms time intervals, respectively, **Fig. 3b, c**). In contrast, the steady-state component (*A*_SS_), which reflects the plateau, was independent of the inter-stimulus interval (**Fig. 3d)**. Only the kinetics of the rapid component changed below 300 ms inter-stimulus interval (rR: 5.7 ± 0.4 ms to 3.3 ± 0.2 ms from 40 to 300 ms time intervals, respectively, **Fig. 3e**). Although the amplitude of the short-term adaptation changed with the inter-stimulus interval, the short-term time constant (τ*_ST_*) was unaffected (**Fig. 3f)**. Finally, the PSTR peak-to-plateau ratio was stable for silent intervals longer than 200 ms (4.7 ± 0.2; **Fig. 3g**). Together, these results are consistent with forward-masking experiments, in which no masking occurs when the time interval between the masker and the signal equals or exceeds 200 ms (Harris and Dallos, 1979; Relkin and Doucet, 1991). Consequently, we decided to carry out all the experiments using a 300 ms inter-stimulus time interval.

**Fig. 3.**
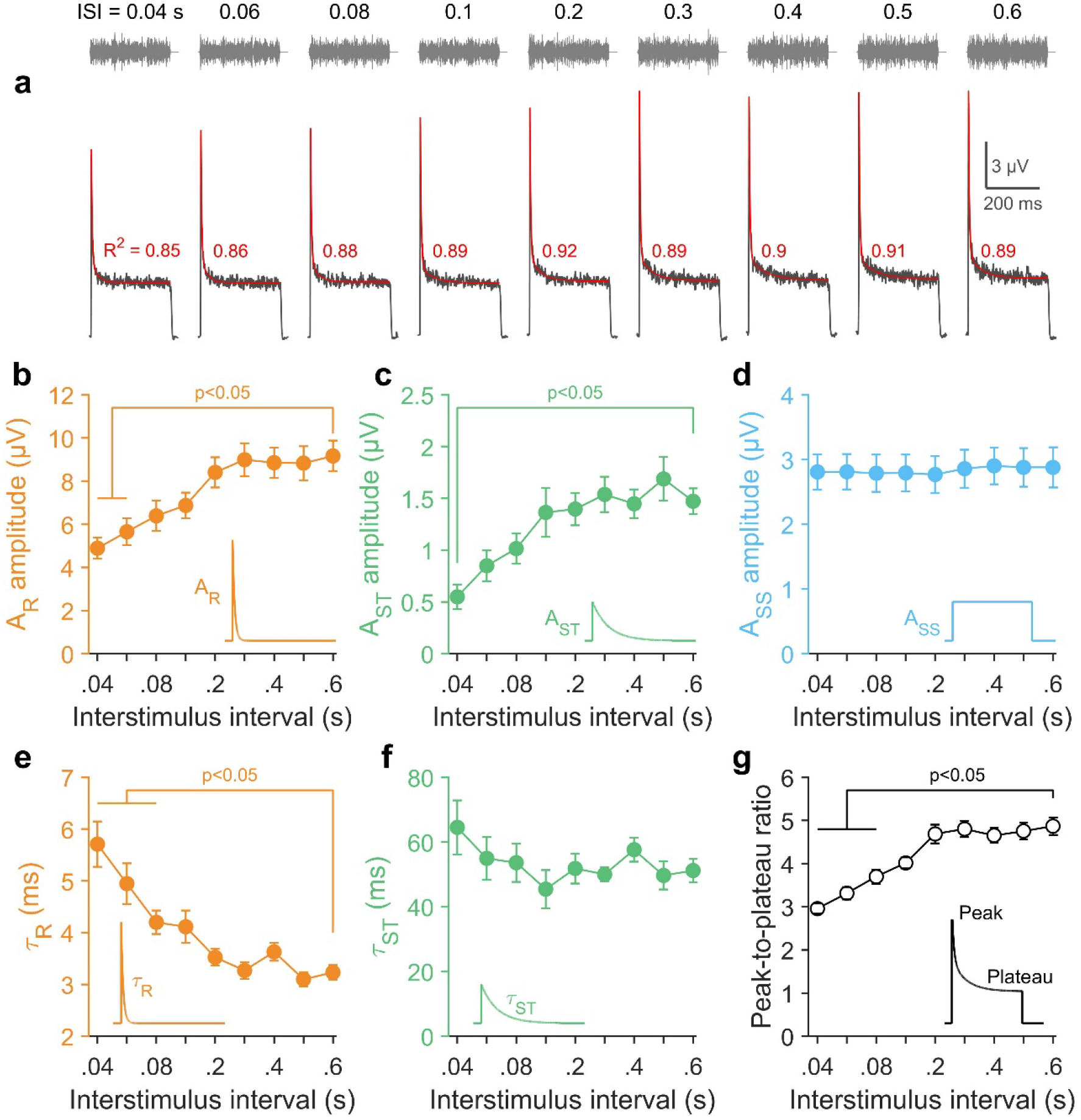
PSTR as a function of the inter-stimulus time interval (ISI). **a** PSTRs (bottom traces) in response to a burst of a noise band centered at 4 kHz (top traces) and varying in inter-stimulus interval (ISI) from 0.04 to 0.6 s (data from one representative gerbil). The level of presentation was fixed at 50 dB SPL. In red, the fitted model (*i*.*e*. two decaying-exponential components plus a constant term) and its coefficient of determination (*R*^2^, red digit). **b-d** Amplitude of the three components of the fitting model as a function of the ISI. **e, f** Time constant of the rapid and short-term components as a function of the ISI. **g** PSTR peak-to-plateau ratio as a function of the ISI. Data are mean ± sem, *n* = 12 gerbils. Statistical comparisons between pairs of samples were performed using ISI = 0.6 s as the reference.

#### Simultaneous recordings of peri-stimulus time response and single units

To investigate the relationship between the PSTHs of single ANFs and the PSTR recorded at the round window, we carried out concomitant measurements using a sharp glass pipette in the gerbil auditory nerve and a silver ball electrode in the round window niche, respectively. For each ANF, the SR and the characteristic frequency (CF) were determined (**Fig. 4a**). Like the PSTR, the fiber was then stimulated with a 1/3 octave noise band centered on its CF. Here, we collected 43 simultaneous recordings.

**Fig. 4.**
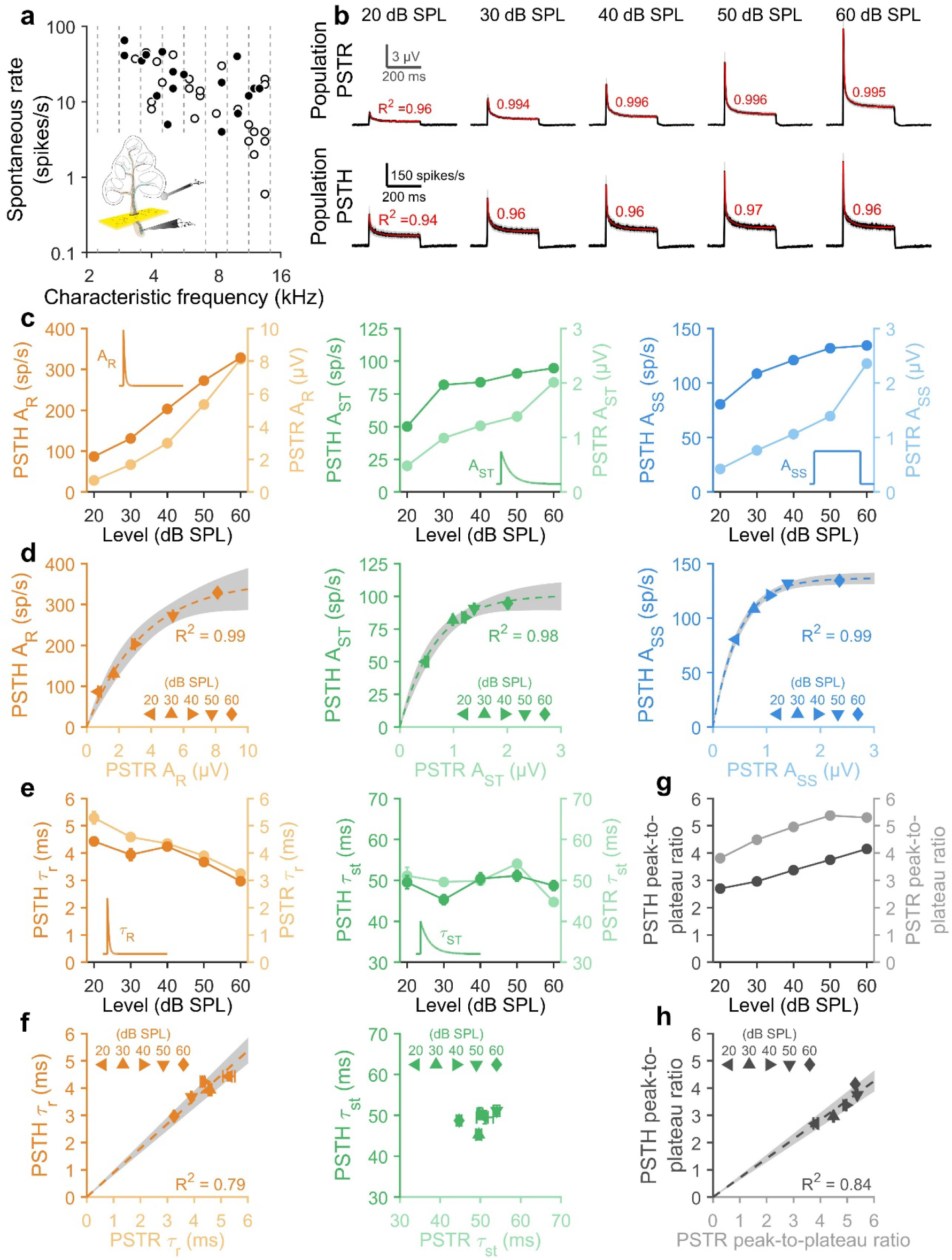
Paired recordings in response to increasing sound level. **a** Spontaneous rate and characteristics frequency of the fibers used in this analysis. For 17 of the 43 fibers (shown with closed symbols), paired recordings were successfully performed from 20 to 60 dB SPL, in 10 dB steps. **b** Mean PSTR (top traces) and mean PSTH (bottom traces) obtained in response to a burst of 1/3 octave band noise centered on the fiber’s CF, varying in level from 20 to 60 dB SPL (*n* = 17 ANFs). In red is shown the fitting model with its coefficient of determination (R^2^, red digit). **c** Amplitude parameters derived from population PSTR (dark color, left axis) and PSTH (light color, right axis), as a function of the sound level. **d** Correlation between amplitude parameters derived from population PSTH (*y*-axis) and population PSTR (*x*-axis), for the different sound levels shown above in panel C. Dashed line is a fit to the data using an exponential model (*A*_R_: *y*=358×(1-exp(-*x*/3.5)), *A*_ST_: *y*=101× (1-exp(-*x*/0.65)), *A*_SS_: *y*=137× (1-exp(-*x*/0.48))). **e** Time-constant parameters derived from population PSTR (dark color, left axis) and PSTH (light color, right axis) as a function of the sound level. **f** Correlation between time-constant parameters derived from population PSTH (*y*-axis) and population PSTR (*x*-axis) for the different sound levels shown above in panel E. In F, the dashed line is a fit to the data using a linear model (*y*=0.9×*x*). **g** Peak-to-plateau ratio derived from population PSTR (dark color, left axis) and PSTH (light color, right axis), as a function of the sound level. **h** Correlation between peak-to-plateau ratio derived from population PSTH (*y*-axis) and population PSTR (*x*-axis) for the different sound levels shown above in panel g. Dashed line is a fit to the data using a linear model (*y* = 0.71×*x*). In panels **d, e, f**, and **h**, the gray area gives the predicted range of the fit for a confidence level at 95%.

#### Dependence on sound pressure level

First, we correlated the activity of several ANFs to the corresponding PSTR, comparing the amplitudes and time constants in both PSTR and PSTH populations as a function of sound level. To do so, we selected 17 fibers for which a complete experimental protocol was available, *i*.*e*., successful recording of activity from 20 to 60 dB SPL (**Fig. 4a**, filled symbols). Although both PSTR and PSTH populations behaved similarly (**Fig. 4b; Supp. Fig. 1a**), amplitude-fitting parameters derived from PSTHs tended to saturate for supra-threshold levels (especially for *A*_ST_ and *A*_SS_), whereas those of PSTRs increased more slowly and did not saturate **(Fig. 4c; Supp. Fig. 1b-d**), probably because of cochlear spread of excitation. Plotting the amplitude-fitting parameters of PSTR (*A*_R_, *A*_ST_ and *A*_SS_) against those of PSTH displayed an exponential relationship (**Fig. 4d**). The rapid time constant τ*_R_* decreased with increasing sound level, but the short-term time constant τ*_st_* was insensitive to sound level (**Fig. 4e; Supp. Fig. 1e, f**). Although the PSTH τ*_R_* was slightly shorter at the lowest levels (*i*.*e*., 20 and 30 dB SPL), we found a significant correlation (*R*^2^ = 0.79, *p* = 0.023) with the PSTR τ*_R_* populations (**Fig. 4f**). Because the PSTH τ*_ST._*was independent of the stimulation level around 50 ms duration, no significant correlation was found (**Fig. 4f**). Finally, PSTR peak-to-plateau ratios were larger, but significantly correlated (*R*^2^ = 0.84, *p* = 0.025) with those of the PSTH (**Fig. 4g, h; Supp. Fig. 1g**).

#### Dependence on probe frequency

In the gerbil, IHCs from the apical half of the cochlea are mainly innervated by high-SR ANFs, whereas basal IHCs are innervated by ANFs with a greater SR diversity (Schmiedt, 1989; Ohlemiller and Echteler, 1990; Müller, 1996; Bourien et al., 2014; Huet et al., 2016, 2019). When stimulated at a fixed intensity above their threshold, the rapid adaptation kinetics are faster for high-SR fibers (*i*.*e*., those in the majority in the apex) than for low-SR fibers. In addition, the high-SR fibers exhibit a larger PSTH peak-to-plateau ratio than the low SR-fibers (Westerman and Smith, 1984; Rhode and Smith, 1985; Müller and Robertson, 1991; Relkin and Doucet, 1991). Assuming that the PSTR reflects the sum of PSTHs derived from single units, we used our paired data to compare amplitudes and time constants in response to stimuli presented at 40 dB above PSTR thresholds. The 43 simultaneous recordings (see **Fig. 4a**) were pooled per 1/3 octave frequency bandwidths according the fiber’s CF (3.15 to 12.6 kHz, 1/3 octave steps; **Fig. 5a, b**). The shape of averaged PSTRs and PSTHs behaved similarly across frequencies, with a larger peak-to-plateau ratio and a shorter τ*_R_* for low probe frequencies (**Fig. 5a, b)**. Indeed, a significant linear correlation (*R* = 0.99, *p* < 0.001) was observed between PSTH and PSTR rR. The rapid time constant τ*_R_* was almost the same using both techniques (*y* ::: 0.95 × *x*, **Fig. 5c**). We then investigated the relationship between the PSTR peak-to-plateau ratio and the mean SR in each CF-based pool of ANFs. Here again, a significant correlation (*R* = 0.97, *p* < 0.001) was found. The larger the PSTR peak-to-plateau ratio, the higher was the mean SR of fiber populations *y* ::: 6.1 × *x* - 7.4, **Fig. 5d**).

**Fig. 5.**
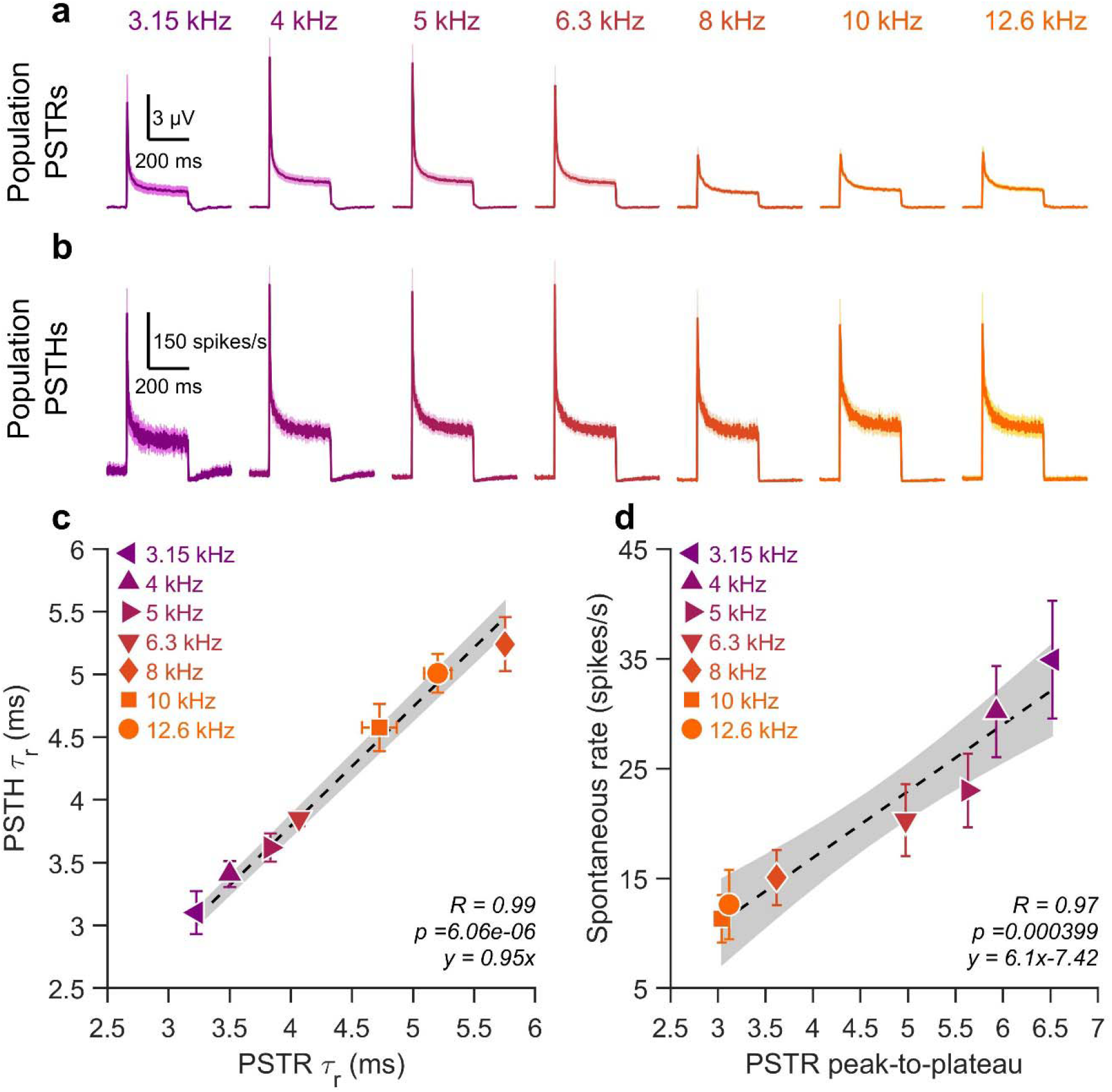
Paired recordings in response to different probe frequencies. **a, b** Population PSTRs (**a**) and PSTHs (**b**) obtained per 1/3 octave band around the fiber’s CF, in response to a 50 dB SPL stimulation presented at the fiber’s CF. 43 ANFs were considered in this analysis (see Fig. 4a), with a minimum of 3 fibers per 1/3 octave band. **c** Rapid time constant derived from PSTH (*y*-axis) as a function of those derived with PSTR (*x*-axis) for different CF (3.15 in purple to 12.6 kHz in orange). Dashed line is a linear fit to the data (*y* = 0.95×*x*). **d** Spontaneous rate of the fibers per 1/3 octave band (*y*-axis) as a function of PSTR peak-to-plateau (*x*-axis) for different probe frequencies varying from 3.15 (purple) to 12.6 kHz (orange). The fibers were pooled per 1/3 octave band according to their CF. In panels **c** and **d**, the gray area gives the predicted range of the fit for a confidence level of 95%.

### Predictive modelling

#### Validation of the predictive model in gerbils

For both PSTH τ*_R_* and ANF’s SR, a prediction interval with a confidence level of 95% was obtained from linear fits (gray area in **Fig. 5c, d**). The strength of the prediction was then tested on two independent groups of gerbils. In the first group, the PSTRs were only measured at 50 dB SPL (*i*.*e*. 40 dB above the PSTR threshold) (**Fig. 6a; Supp. Fig. 2a**). In the second group, only single-unit recordings were performed (dataset published as Huet et al, 2016 (Huet et al., 2016)) to extract the fiber’s SR and PSTH measured at 40 dB above the PSTR threshold. When measuring PSTR alone, the *A*_R_ dramatically decreased with frequency (8.1 ± 0.9 to 2.3 ± 0.3 µV from 3.15 to 16 kHz, respectively**; Supp. Fig. 2b**) and τ*_R_* increased (from 3.6 ± 0.2 to 4.3 ± 0.3 ms from 3.15 to 16 kHz, respectively; **Supp. Fig. 2e**) with a cut-off frequency at 6.3 kHz. No significant change was observed for the short-term and steady-state components (**Supp. Fig. 2, c, d, f**). Together, this led to a reduction of the PSTR peak-to-plateau ratio (6.2 ± 0.5 to 3.5 ± 0.2 from 3.15 to 16 kHz, respectively; **Supp. Fig. 2g**). When plotting PSTH τ*_R_* (**Fig. 6b**) measured from single ANFs pooled per 1/3 octave band according their CFs, they all fall within the 95% prediction interval (*gray area*) derived from the PSTR (**Fig. 6c**). Also, the measured PSTH τ*_R_* of ANFs and the PSTR-based prediction showed a high degree of correlation (*R* = 0.98, see inset **Fig. 6c**).The mean SR measured from the same ANF populations also matched the prediction interval derived from the PSTR peak-to-plateau ratio remarkably well (**Fig. 6d**), with a strong correlation coefficient (*R* = 0.94, **Fig. 6e**).

**Fig. 6.**
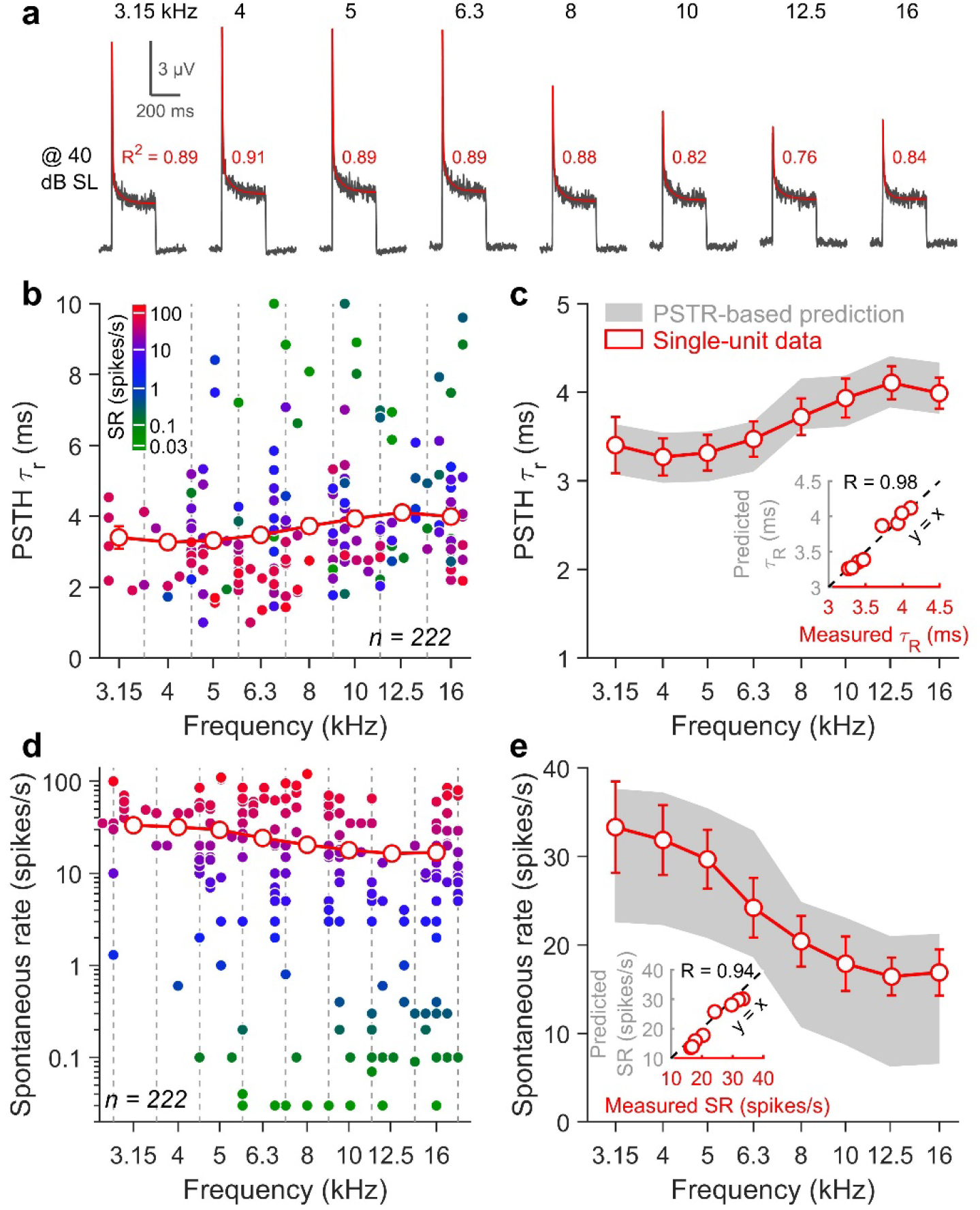
Prediction of fiber PSTH τ_R_ and SR in gerbils. **a** PSTRs obtained in response to a burst of 1/3 octave band of noise centered at a probe frequency varying from 3.15 to 16 kHz (data obtained in one representative gerbil). In red is shown the fitted model (*i*.*e*. two decaying-exponential components plus a constant term) and its coefficient of determination (R^2^, red digit). **b** Scatter plot of PSTH τ_R_ as a function of the fiber’s CF for 222 ANFs (closed symbols). The fiber’s SR is represented using a color scale from green (low-SR) to red (high-SR fiber). The red open symbols show the mean PSTH τ_R_ (± 1 sem) when fibers were pooled per 1/3 octave band according to CF. **c** Same data as in panel B but without individual data and using a *y*-axis magnification. Gray area gives a PSTR-based prediction range of the mean PSTH τ_R_ using the predictive model shown in Fig. 5c. **d** Scatter plot of fiber SR as a function of CF for 222 ANFs (closed symbols). The red open symbols show the mean SR (± 1 sem) when fibers were pooled per 1/3 octave band according to CF. **e** Same data as in panel **d**, but without individual data and using a linear *y*-axis scale. Gray area gives a PSTR-based prediction range of the mean SR using the predictive model shown in Fig. 5d. Note the high degree of correlation between predicted and measured PSTH τ_R_ (*R* = 0.98) and SR (*R* = 0.94) values.

#### Validation of the predictive model in mice

To validate the PSTR as a predictive tool for the functional state of ANFs in another animal model, we recorded PSTRs in C57BL/6 mice using probe frequency varying from 5.7 to 32 kHz in 1/2 octave steps (**Fig. 7a; Supp. Fig. 3a)**. In contrast to the gerbils, the amplitude of the rapid components (**Supp. Fig. 3b)** was independent of the probe frequency, as well as the short-term and the steady-state components **(Supp. Fig. 3c, d)**. However, the rapid time constant τ*_R_* increases with the stimulation frequency (3.1 ± 0.4 ms to 4.1 ± 0.3 ms for 5.7 and 32 kHz, respectively; **Supp. Fig. 3e**). As also shown in gerbils, τ*_ST_* was independent of the probe frequency (40.6 ± 2.8 ms across frequency, **Supp. Fig. 3f**). Together with the lack of change in the PSTR peak-to-plateau ratio (**Supp. Fig. 3g**), these PSTR characteristics of the mice suggests a more homogeneous distribution of ANFs along the tonotopic axis than in gerbils. To validate this hypothesis, we re-analyzed the single-unit recordings in the auditory nerve of mice as published by Taberner and Liberman in 2005(Taberner and Liberman, 2005). Due to the small number of recordings in C57BL/6 (58 ANFs), and the lack of striking differences to CBA/CaJ mice, we only reported results from CBA/CaJ mice. Among the 196 ANFs, we analyzed the 130 units stimulated at 40 dB above PSTR thresholds. When fibers were pooled per 1/2 octave band according their CFs, the mean PSTH τ*_R_* increased with frequency (**Fig. 7b**) as predicted by the PSTR τ*_R_* **(Fig. 7c**, *R* = 0.97). Also, the mean fiber SR calculated from the same 130 fibers remained stable across frequency (**Fig. 7d**) and fell within the 95% prediction interval (*gray area*) derived from the PSTR peak-to-plateau ratio (**Fig. 7e**, *R* = 0.62).

**Fig. 7.**
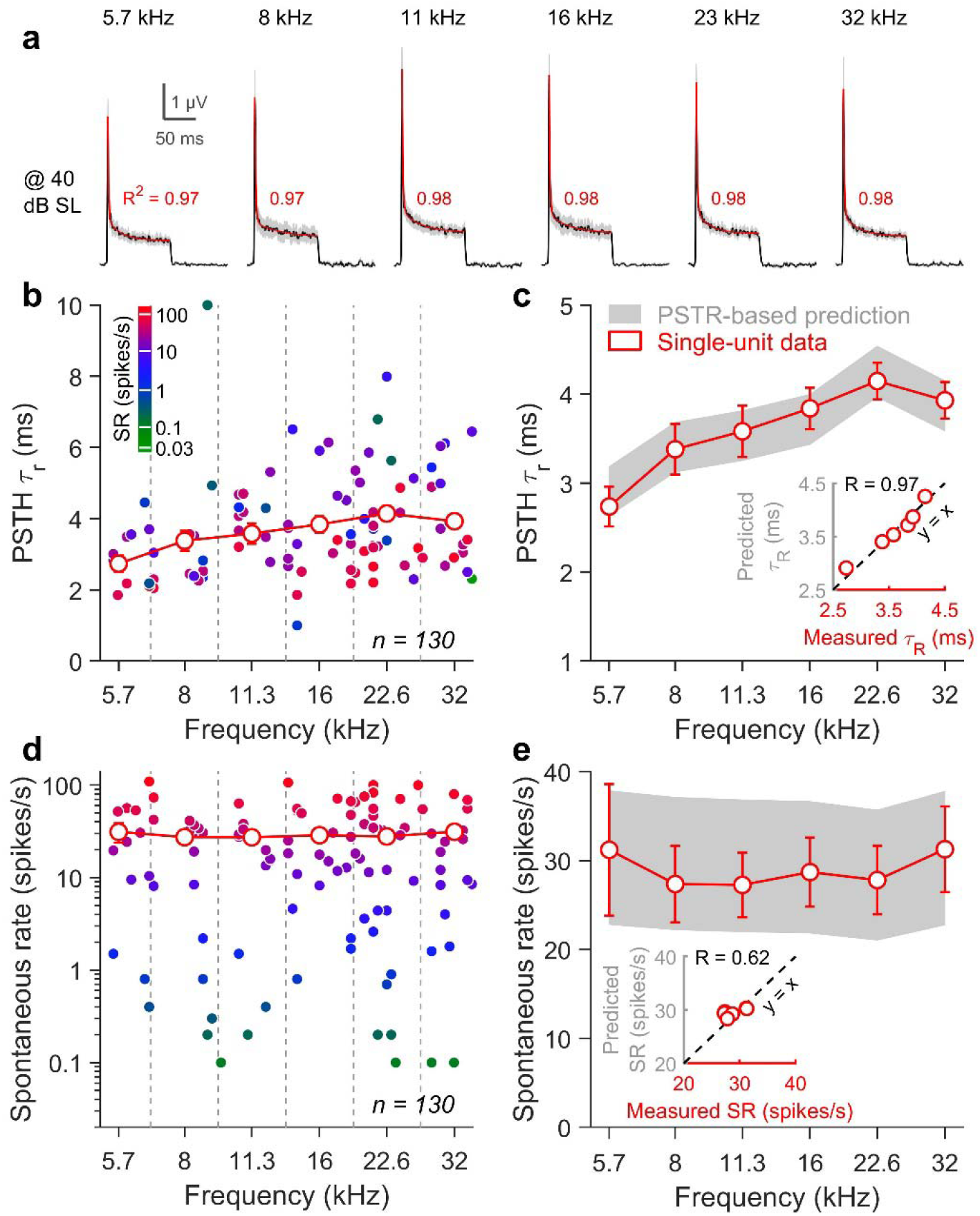
Prediction of fiber PSTH τ_R_ and SR in mice. **a** PSTRs obtained in response to a burst a 1/3 octave noise band centered on a probe frequency varying from 5.7 to 32 kHz. In red is shown the fitting model (*i*.*e*. two decaying-exponential components plus a constant term) and its coefficient of determination (R^2^, red digit). **b** Scatter plot of PSTH τ_R_ as a function of the fiber’s CF for 130 ANFs (closed symbols). The fiber’s SR is represented using a color scale from green (low-SR) to red (high-SR fiber). The red open symbols show the mean PSTH τ_R_ (± 1 sem) when fibers were pooled per 1/3 octave frequency band according to CF. **c** Same data as in panel B without individual data and using a *y*-axis magnification. Gray area gives a PSTR-based prediction range of the mean PSTH τ_R_ using the predictive model shown in Fig. 5c. **d** Scatter plot of fiber’s SR as a function of the fiber’s CF for 130 ANFs (closed symbols). The red open symbols show the mean SR (± 1 sem) when fibers were pooled per 1/3 octave band according to CF. **e** Same data than in panel **d** without individual data and using a linear *y*-axis scale. The gray area gives a PSTR-based prediction range of the mean SR using the predictive model shown in Fig. 5d. Note the correlation between the predicted and measured PSTH τ_R_ (*R* = 0.97). Although the correlation index was lower (*R* = 0.62), the SR values are distributed along the *y* = *x* line.

#### Proof of concept in humans

For obvious ethical reasons, single unit recordings from the cochlear nerve have never been performed in humans, creating a critical gap in our understanding of the applicability of data from animal models. We took advantage of cerebellopontine angle surgeries to record PSTRs from an electrode placed on the intracranial portion of the cochlear nerve in consented patients with normal hearing (**Supp. Fig. 4**). Those patients were undergoing neurosurgeries for cranial nerve functional disorders (trigeminal neuralgia and hemifacial spasm (Møller and Jannetta, 1981; Møller and Jho, 1989)). Click-evoked CAPs were recorded intraoperatively to ensure that no noticeable changes occurred in auditory nerve function as a result of the surgical dissection (**Fig. 8a)**. Then, PSTR were recorded in response to 1/3 octave band noise centered on 4 kHz, presented 40 dB above the click-evoked CAP threshold. The waveform of the human PSTR was similar to those recorded from gerbils or mice (**Fig. 8b**). The amplitude of the rapid (*A*_R_: 2.0 ± 0.3 µV), short-term (*A*_ST_: 0.1 ± 0.1 µV) and steady state (*A*_*S*S_: 0.75 ± 0.08 µV) components were in the range of those measured in rodents. The rapid time constant τ*_R_* was equal to 2.8 ± 0.07 ms, which is in the same range as the animal data **(Fig. 8c)**. The PSTR peak-to-plateau ratio value was 5.0 ± 0.1, predicting a mean SR of 23 ± 3 spikes/s in the 4 kHz region of the human auditory nerve (**Fig. 8d**). Together, our results suggest that the PSTR constitutes a promising tool to extract the kinetics of neural adaptation to sound stimulation and to map the SR-based composition of the ANFs in humans and other mammalian species.

**Fig. 8.**
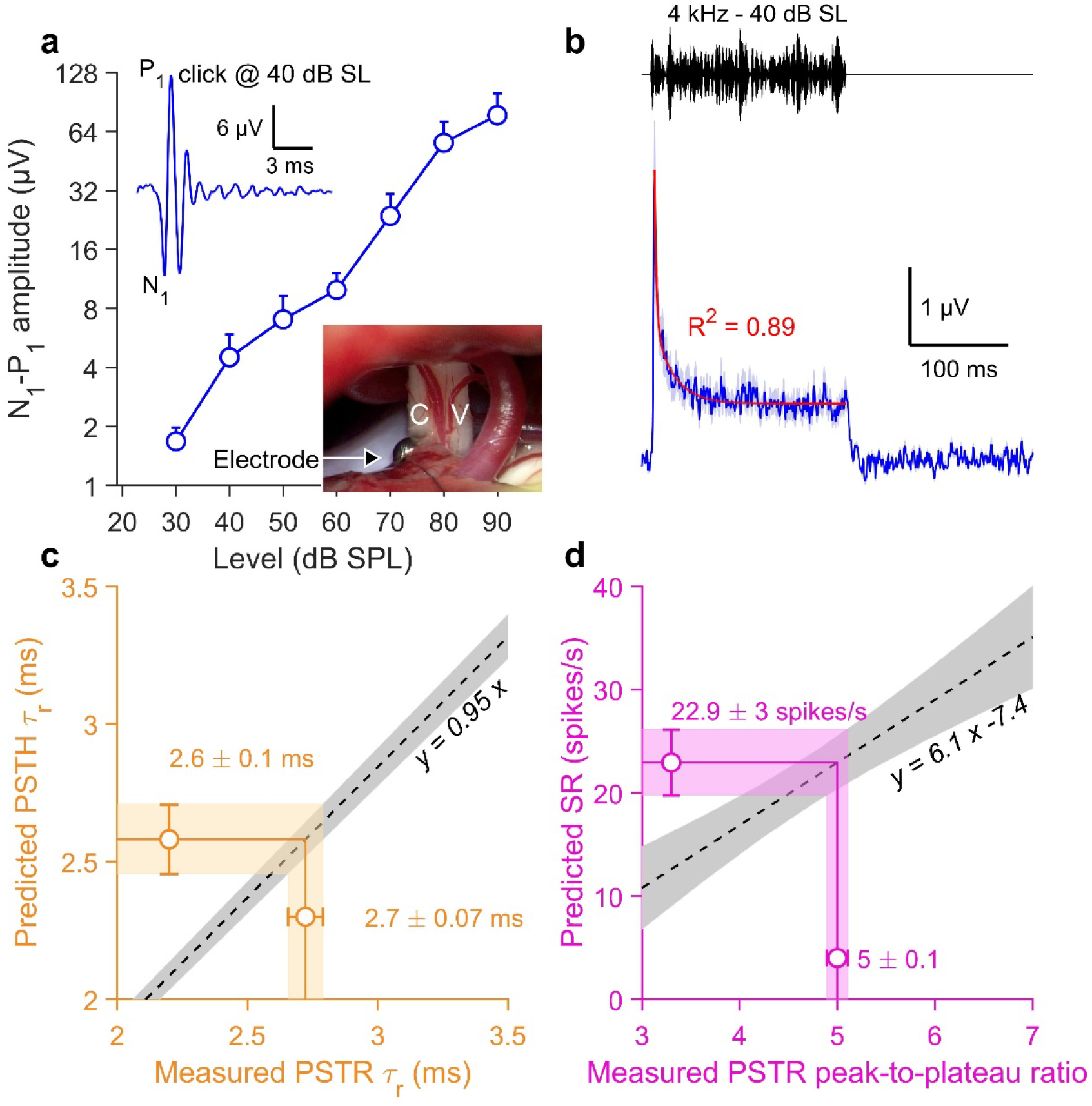
Prediction of fiber PSTH τ_R_ and SR at 4 kHz in humans. **a** CAP N_1_-P_1_ amplitude (mean ± sem) in response to clicks as a function of the sound level. A threshold at 30 dB SPL together with a double slope in the growth function attest to normal hearing. *Top inset*: Example of a CAP in one representative patient (200 presentations, same polarity, 40 dB above threshold). *Bottom inset*. Auditory nerve responses were measured using a ball electrode directly on the surface of the eighth nerve (C, cochlear nerve; V, vestibular nerve). **b** PSTR to a 200-ms burst of 1/3 octave noise band centered at 4 kHz and presented at 40 dB above the click-evoked CAP threshold (100 presentations). The blue solid line indicates the mean PSTR across the 8 patients, ± 1sem (light blue area). In red is shown the fitted model (*i*.*e*. two decaying-exponential components plus a constant term) and its coefficient of determination (*R*^2^, red digit). **c** Prediction of the PSTH τ_R_ (vertical error bar) using the measured PSTR τ_R_ at 40 dB above click-evoked CAP threshold (horizontal error bar) and the predictive model identified in gerbils (gray area). **d** Prediction of the mean SR (vertical error bar) using the measured PSTR peak-to-plateau ratio at 40 dB above click-evoked CAP threshold (horizontal error bar) and the predictive model identified in gerbils (gray area).

## Discussion

Using far-field recordings, we probed the adaptation kinetics and spontaneous action potential firing in limited populations of ANFs in rodents. In addition, we examined whether the relationship between the PSTR and PSTH allows the prediction of these two electrophysiological features in normal-hearing human ANFs.

### Adaptation of auditory nerve fibers

In response to the onset of an acoustic stimulus, the spike rate of an ANF rapidly increases to a maximum value and thereafter adapts to reach a steady-state rate. The post-onset reduction in discharge rate has been attributed mainly to the reduction of vesicle release at the IHCs ribbon synapse (Beutner et al., 2001; Goutman and Glowatzki, 2007). Initial exocytosis operates at a high rate at the onset of stimulation, but slows within a few milliseconds, as the readily releasable pool is depleted. Other mechanisms, including the desensitization of post-synaptic glutamate receptors and refractoriness of action potential generation at the ANF level, may also contribute (Heil and Peterson, 2015). The most definitive experimental procedure to investigate ANF adaptation relies on single-unit recording. However, this approach is very invasive and difficult to achieve, as the auditory nerve is deep to the cerebellum and largely surrounded by the petrous portion of the temporal bone. Here, we investigated whether far-field PSTR recorded at the round window replicates single-unit PSTH, especially with respect to the rapid and short-term time constants of the post-onset adaptation. As in prior ANF studies (Westerman and Smith, 1984), we used a double exponential function to fit the PSTR waveform to the PSTH analysis. With increasing sound level, we saw a reduction of the rapid time constant. Consistently, Westerman and Smith reported a decrease of the PSTH rapid time constant from 10 to 1 ms as tone level increased (Westerman and Smith, 1984), which is in the range our PSTR data. In addition, the short-term time constant was independent of the stimulus level, and approximates 50 ms, which is also in agreement with our PSTR measurements To most directly evaluate how a population of ANFs contributes to the PSTR, we carried out simultaneous recordings of single ANFs and round-window PSTRs. Pair-by-pair analysis showed many similarities, such as the shape and the time constants whatever the frequency and the level of sound stimulation. Few discrepancies are however inherent to the nature of the recording techniques. For example, PSTH amplitudes (especially *A*_ST_ and *A*_SS_) tend to saturate at supra-threshold levels, while those of PSTR continue to increase. Because PSTHs reflect the time course of the discharge rate in response to sound, their amplitudes are limited by the maximum discharge rate of the fiber, and the relatively limited dynamic range of single ANFs. In contrast, the PSTR amplitude can continue to grow above 50 dB SPL, because of the spread of excitation along the organ of Corti, which recruits ANFs with characteristic frequencies outside the nominal bandwidth of the noise. Thus, at least within a range of 40 dB above threshold, PSTRs may accurately reflect the rapid and short-term adaptation of the single ANFs with CFs in the noise bandwidth. We further reported larger peak-to-plateau ratio for PSTRs than for PSTHs. This difference is due to the choice of the bin size to build the PSTH (*i*.*e*. 0.5 ms in the present study). If the bin size is too large, adaptation cannot be measured. If it is too small, the time histogram fluctuates rapidly and the underlying spike rate cannot be discerned. On the other hand, because PSTR is an electrical signal recorded in the far field, the peak-to-plateau ratio depends on the temporal resolution of the acquisition system. Nevertheless, the similarly rapid time constants measured with both techniques makes the PSTR a powerful method to evaluate fast adaptation of the ANFs in animal models in which single unit recordings are difficult to achieve.

### Spontaneous activity of auditory nerve fibers

Several studies have described the SR patterns of ANFs in the gerbil cochlea, with a majority of high-SR fibers in the apical part and a more balanced distribution of high-, medium- and low-SR fibers in the basal half (Schmiedt, 1989; Ohlemiller and Echteler, 1990; Müller, 1996; Huet et al., 2016). This specificity makes the gerbil a unique model to investigate the function of different pools of ANFs (Bourien et al., 2014). High-SR fibers exhibit greater peak-to-plateau ratios than low-SR fibers (Westerman and Smith, 1984). This is also true in cats (Rhode and Smith, 1985), guinea pigs (Müller and Robertson, 1991), and mice (Taberner and Liberman, 2005). Because the apical cochlea in gerbil is mainly populated by high-SR fibers, the peak-to-plateau ratio should be larger for PSTRs in response to lower frequency sounds, as we indeed observed. Thus, the PSTR peak-to-plateau ratio may be used to infer the mix of ANF SRs at different frequency regions.

Direct demonstration of PSTR-based predictions comes from our paired recordings in gerbil. We saw a strong correlation between the PSTR and PSTH rapid adaptation τ*_R_*. For both parameters, the rapid time constant increases with frequency, which fits well with our previous single-unit measurements (Huet et al., 2016). We also report a decrease in the SR of fiber populations from ∼33 spikes/s at 3.1 kHz to ∼17 spikes/s at 16 kHz. Then, we investigated the predictive value of PSTR in inferring the rapid adaptation time constants and the SRs of fiber populations in another species. When recorded on the cochlear round window of mice, the PSTR peak-to-plateau ratio is mostly invariant across frequency, which is consistent with a homogeneous distribution of fibers along the tonotopic axis of the mouse cochlea. In addition, the mean SR of fiber populations measured in mice (∼29 spikes/s from 5.7 to 32 kHz), matches our prediction from PSTR. We thus conclude that far-field PSTRs constitute a powerful and less invasive tool to rapidly extract key features of the spontaneous and sound-evoked responses of the auditory nerve in animal models, including mice, in which single-unit recordings are difficult to achieve.

### Toward a diagnostic tool

To date, the only functional data on the human ANFs come from CAP recordings, which only reflect the synchronized activity at stimulus onset (see Eggermont, 2017 (Eggermont, 2017) for a review). For ethical reasons (mainly the need to penetrate the auditory nerve with microelectrodes), single-unit recording from the auditory nerve is not feasible in humans, making the composition of the nerve impossible to investigate. Here, we took advantage of cerebellopontine angle surgeries to record PSTRs from an electrode placed on the intracranial portion of the cochlear nerve in patients with normal or sub-normal hearing that were undergoing microvascular decompression (as already described by Møller and Jannetta in 1981(Møller and Jannetta, 1981)). PSTR evoked by noise bands centered on 4 kHz had a similar shape to those recorded in animals. The rapid adaptation of a few milliseconds (∼ 3ms) is in the range of those observed in animals. Based on the PSTR peak-to-plateau ratio, we predict a mean fiber SR of 23 spikes/s in the 4 kHz region, providing a glimpse of the adaptation and the mean spontaneous activity of ANFs in this area of the human cochlea.

Some technical limitations must, however, be taken into account before definitive validation of human data. First, the electrical and acoustic noise levels in the operating room were larger than they are in an experimental laboratory. Second, due to time limitations, we did not record as large range of sound intensities and frequencies as we did in animals. Third, we cannot be sure that anaesthesia or the surgical manipulations needed to expose the eighth nerve may not have changed the physiology of the ANFs. To collect more data, a non-invasive technique is thus need, especially to record PSTRs in awake subjects. Given that the spontaneous neural noise (900 Hz peak) can be extracted in the human using ear canal ECochG recording techniques (Pardo-Jadue et al., 2017), we expect that the signal-to-noise ratio is sufficient to extract neurophonic responses using a tympanic electrode. If not, we can alternatively use the classical transtympanic ECochG. To further investigate the weighted contribution of each pool of ANFs to the PSTR, and to predict their behaviour under pathological conditions such as such ANF loss (hidden hearing loss (Kujawa and Liberman, 2009; Wu et al., 2019)) and/or hair cell loss (deafness (Fernandez et al., 2020; Wu et al., 2020)), we need to develop a mathematical model of the human cochlea. This should be based on the morphological observations (Spoendlin and Schrott, 1989, 1990; Wu et al., 2020), as done previously for the guinea-pig cochlea (Bourien et al., 2014). In the end, we expect the PSTRs to be a powerful diagnostic tool to capture information on auditory nerve survival and, importantly, SR-based function and dysfunction in humans, providing a better understanding of neuropathies, tinnitus, and hyperacusis.

## Acknowledgments

The authors gratefully thank Prof. André Chays and Dr. Arnaud Bazin who initiate the clinical study. We also acknowledge Arthur Lemolton for its technical help. The authors acknowledge language services (www.stels-ol.de) for editing assistance

## Funding

This work was supported by Agence Nationale pour la Recherche (ANR-13-JSV1-0009-01), Inserm Grant (U1051-Dot 02±2014), Cochlear France Award (R11055FF/RVF11006FFA), Gueules Cassées (R20113FF), Fondation Pour l’Audition (FPA RD-2016-2) and the National Institute on Deafness and other Communication Disorders (R01 DC0188).

## Competing interests

The authors declare no competing interests.

## Notes

### Competing Interest Statement

The authors have declared no competing interest.

## References

Batrel C, Huet A, Hasselmann F, Wang J, Desmadryl G, Nouvian R, Puel J-L, Bourien J (2017) Mass Potentials Recorded at the Round Window Enable the Detection of Low Spontaneous Rate Fibers in Gerbil Auditory Nerve. PloS One 12:e0169890.

Beutner D, Voets T, Neher E, Moser T (2001) Calcium dependence of exocytosis and endocytosis at the cochlear inner hair cell afferent synapse. Neuron 29:681–690.

Bourien J, Tang Y, Batrel C, Huet A, Lenoir M, Ladrech S, Desmadryl G, Nouvian R, Puel J-L, Wang J (2014) Contribution of auditory nerve fibers to compound action potential of the auditory nerve. J Neurophysiol 112:1025–1039.

Cazals Y, Huang ZW (1996) Average spectrum of cochlear activity: a possible synchronized firing, its olivo-cochlear feedback and alterations under anesthesia. Hear Res 101:81–92.

Dau T, Verhey J, Kohlrausch A (1999) Intrinsic envelope fluctuations and modulation-detection thresholds for narrow-band noise carriers. J Acoust Soc Am 106:2752–2760.

Eggermont JJ (2017) Ups and Downs in 75 Years of Electrocochleography. Front Syst Neurosci 11:2.

Fernandez KA, Guo D, Micucci S, De Gruttola V, Liberman MC, Kujawa SG (2020) Noise-induced Cochlear Synaptopathy with and Without Sensory Cell Loss. Neuroscience 427:43–57.

Goutman JD, Glowatzki E (2007) Time course and calcium dependence of transmitter release at a single ribbon synapse. Proc Natl Acad Sci U S A 104:16341–16346.

Harris DM, Dallos P (1979) Forward masking of auditory nerve fiber responses. J Neurophysiol 42:1083–1107.

Heil P, Peterson AJ (2015) Basic response properties of auditory nerve fibers: a review. Cell Tissue Res 361:129–158.

Huet A, Batrel C, Tang Y, Desmadryl G, Wang J, Puel J-L, Bourien J (2016) Sound coding in the auditory nerve of gerbils. Hear Res 338:32–39.

Huet A, Batrel C, Wang J, Desmadryl G, Nouvian R, Puel JL, Bourien J (2019) Sound Coding in the Auditory Nerve: From Single Fiber Activity to Cochlear Mass Potentials in Gerbils. Neuroscience 407:83–92.

Huet A, Desmadryl G, Justal T, Nouvian R, Puel J-L, Bourien J (2018) The Interplay Between Spike-Time and Spike-Rate Modes in the Auditory Nerve Encodes Tone-In-Noise Threshold. J Neurosci Off J Soc Neurosci 38:5727–5738.

Joris PX, Schreiner CE, Rees A (2004) Neural processing of amplitude-modulated sounds. Physiol Rev 84:541–577.

Kiang NY (1990) Curious oddments of auditory-nerve studies. Hear Res 49:1–16.

Kujawa SG, Liberman MC (2009) Adding insult to injury: cochlear nerve degeneration after “temporary” noise-induced hearing loss. J Neurosci Off J Soc Neurosci 29:14077–14085.

Meyer AC, Frank T, Khimich D, Hoch G, Riedel D, Chapochnikov NM, Yarin YM, Harke B, Hell SW, Egner A, Moser T (2009) Tuning of synapse number, structure and function in the cochlea. Nat Neurosci 12:444–453.

Møller AR, Jannetta PJ (1981) Compound action potentials recorded intracranially from the auditory nerve in man. Exp Neurol 74:862–874.

Møller AR, Jho HD (1989) Response from the exposed intracranial human auditory nerve to low-frequency tones: basic characteristics. Hear Res 38:163–175.

Müller M (1996) The cochlear place-frequency map of the adult and developing Mongolian gerbil. Hear Res 94:148–156.

Müller M, Robertson D (1991) Relationship between tone burst discharge pattern and spontaneous firing rate of auditory nerve fibres in the guinea pig. Hear Res 57:63–70.

Ohlemiller KK, Echteler SM (1990) Functional correlates of characteristic frequency in single cochlear nerve fibers of the Mongolian gerbil. J Comp Physiol [A] 167:329–338.

Pardo-Jadue J, Dragicevic CD, Bowen M, Delano PH (2017) On the Origin of the 1,000 Hz Peak in the Spectrum of the Human Tympanic Electrical Noise. Front Neurosci 11:395.

Relkin EM, Doucet JR (1991) Recovery from prior stimulation. I: Relationship to spontaneous firing rates of primary auditory neurons. Hear Res 55:215–222.

Rhode WS, Smith PH (1985) Characteristics of tone-pip response patterns in relationship to spontaneous rate in cat auditory nerve fibers. Hear Res 18:159–168.

Ryan A (1976) Hearing sensitivity of the mongolian gerbil, Meriones unguiculatis. J Acoust Soc Am 59:1222–1226.

Schmiedt RA (1989) Spontaneous rates, thresholds and tuning of auditory-nerve fibers in the gerbil: comparisons to cat data. Hear Res 42:23–35.

Spoendlin H, Schrott A (1989) Analysis of the human auditory nerve. Hear Res 43:25–38.

Spoendlin H, Schrott A (1990) Quantitative evaluation of the human cochlear nerve. Acta Oto-Laryngol Suppl 470:61–69; discussion 69-70.

Taberner AM, Liberman MC (2005) Response properties of single auditory nerve fibers in the mouse. J Neurophysiol 93:557–569.

Westerman LA, Smith RL (1984) Rapid and short-term adaptation in auditory nerve responses. Hear Res 15:249–260.

Wu PZ, Liberman LD, Bennett K, de Gruttola V, O’Malley JT, Liberman MC (2019) Primary Neural Degeneration in the Human Cochlea: Evidence for Hidden Hearing Loss in the Aging Ear. Neuroscience 407:8–20.

Wu P-Z, O’Malley JT, de Gruttola V, Liberman MC (2020) Age-Related Hearing Loss Is Dominated by Damage to Inner Ear Sensory Cells, Not the Cellular Battery That Powers Them. J Neurosci Off J Soc Neurosci 40:6357–6366.

