## Supplemental Figures for "Probing adaptation and spontaneous firing in human auditory-nerve fibers with far-field peri-stimulus time responses"

This file contains

**Supp. Fig. 1 PSTR as a function of the sound level**

**Supp. Fig. 2 PSTR as a function of the probe frequency in gerbils**

**Supp. Fig. 3: PSTR as a function of the probe frequency in mice.**

**Supp. Fig. 4 Hearing threshold of the human patients.**

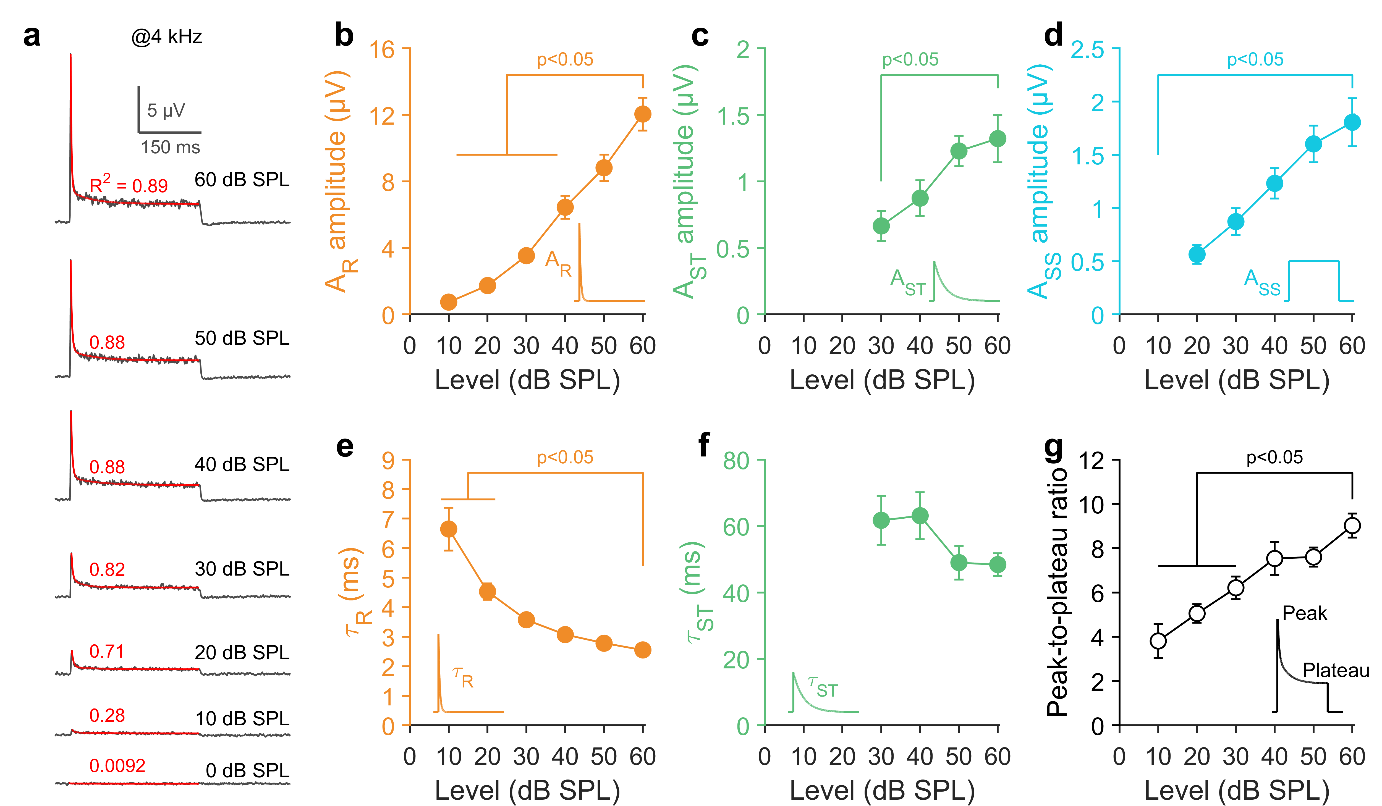

**Supp. Fig. 1 PSTR as a function of the sound level. a** PSTRs obtained in response to a burst of a band of noise centered at 4 kHz and varying in level from 0 to 60 dB SPL (data obtained in one representative gerbil). In red is shown the fitting model (*i.e.* two decaying-exponential components plus a constant term) and its coefficient of determination (R^2^, red digit). **b-d** Amplitude of the three components of the fitting model (see *inset*) as a function of the level. **e, f** Time constants of the rapid and short-term components as a function of the level. **g** The PSTR peak-to-plateau ratio as a function of the level. Data are mean ± sem, *n* = 14 gerbils. Statistical comparisons between pairs of samples were performed using 60 dB SPL as the reference.

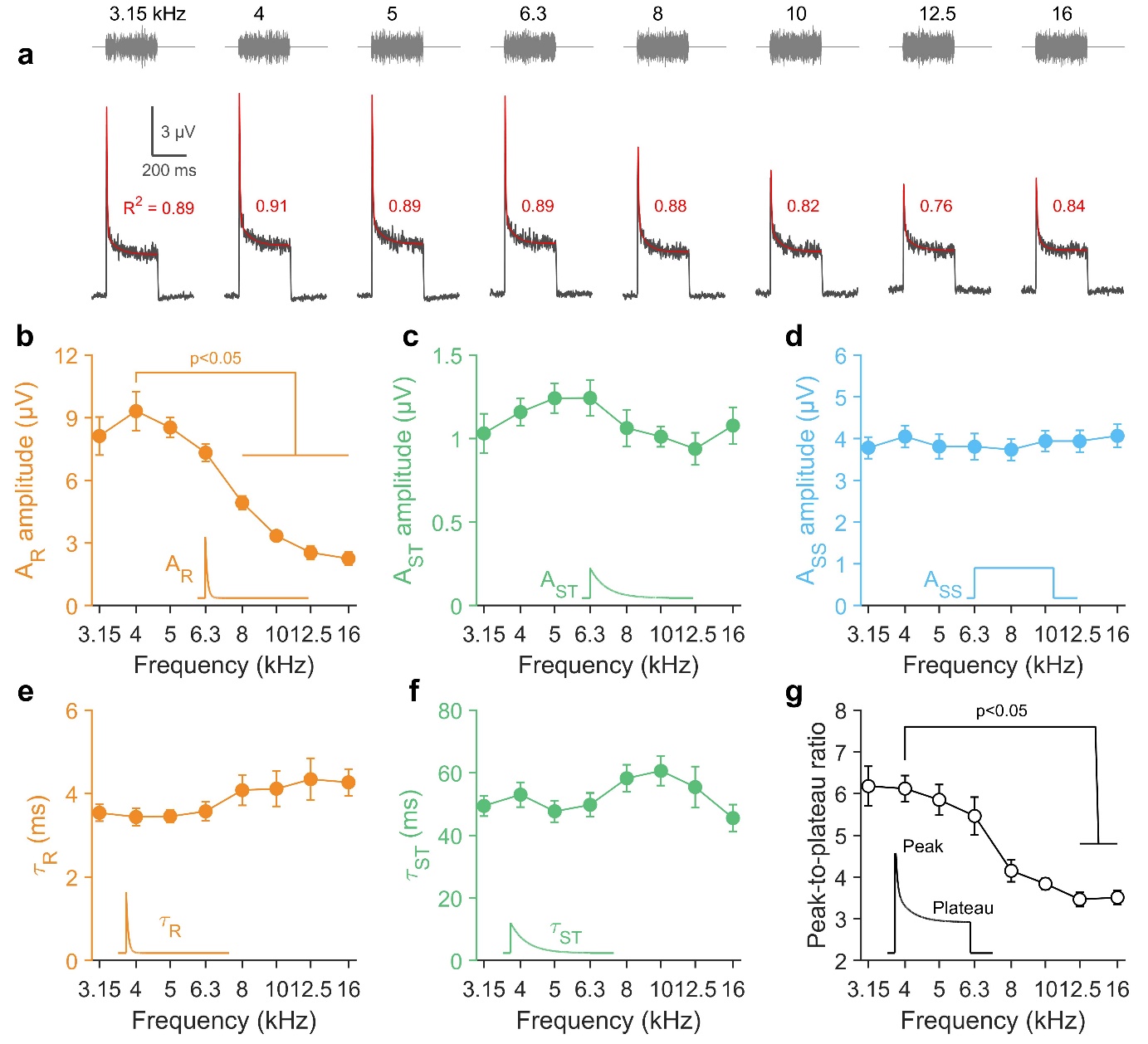

**Supp. Fig. 2 PSTR as a function of the probe frequency in gerbils. a** PSTRs (bottom traces) obtained in response to a burst a 1/3 octave frequency band of noise (top traces) centered to a probe frequency varying from 3.15 to 16 kHz (data obtained in one representative gerbil). In red, the fitting model (*i.e.* two decaying-exponential components plus a constant term) and its coefficient of determination (*R*^2^, red digit). **b-d** Amplitude of three components of the fitting model (see *inset*) as a function of the probe frequency. **e, f** Time constants of the rapid and short-term components as a function of the probe frequency. **g** The PSTR peak-to-plateau ratio as a function of the probe frequency. Data are mean ± sem, *n* = 14 gerbils. Statistical comparisons between pairs of samples were performed using 4 kHz as the reference.

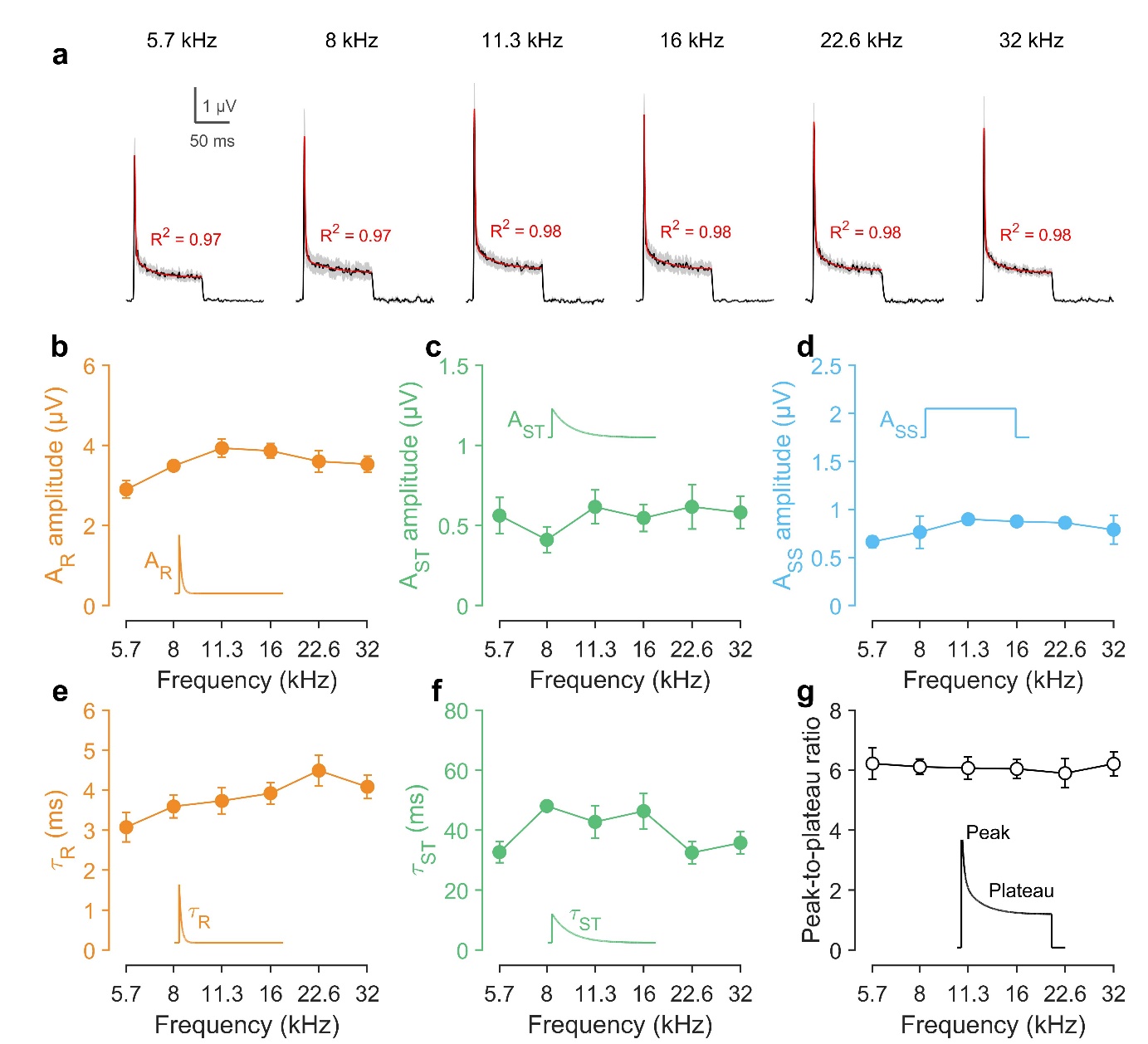

**Supp. Fig. 3: PSTR as a function of the probe frequency in mice.** **a** PSTRs obtained in response to a burst a 1/3 octave frequency band of noise centered to a probe frequency varying from 5.7 to 32 kHz. In red, the fitting model (i.e. two decaying-exponential components plus a constant term) and its coefficient of determination (R^2^, red digit). **b-d** Amplitude of three components of the fitting model (see inset) as a function of the probe frequency. **e, f** Time constants of the rapid and short-term components as a function of the probe frequency. **g** The PSTR peak-to-plateau ratio as a function of the probe frequency. Data are mean ± sem, *n* = 9 mice.

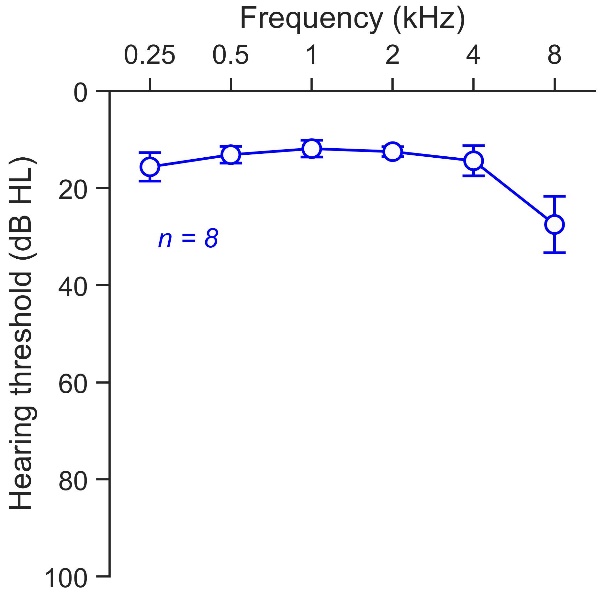

**Supp. Fig. 4 Hearing threshold of the human patients.** These 8 normal-hearing patients (thresholds ≤20 dB HL between 0.5 and 4 kHz) were subject to surgery requiring opening of the cerebellopontine angle (62.1 ± 9.3 years-old).
